# Effects of environmental variables on the distribution of juvenile cubomedusae *Carybdea marsupialis* in the coastal Western Mediterranean

**DOI:** 10.1101/2020.03.10.985515

**Authors:** Cesar Bordehore, Eva S. Fonfría, Cristina Alonso, Beatriz Rubio-Tortosa, Melissa J. Acevedo, Antonio Canepa, Silvia Falcó, Miguel Rodilla, Verónica Fuentes

**Affiliations:** “Ramon Margalef” Environmental Research Institute (IMEM) University of Alicante, San Vicente del Raspeig, Spain; Department of Ecology, University of Alicante, San Vicente del Raspeig, Spain; Institute of Marine Sciences, CSIC, Barcelona, Spain; Polytechnic School, Universidad de Burgos, Burgos, Spain; Research Institute for Integrated Management of Coastal Areas, Universitat Politècnica de València, Gandia, Spain

## Abstract

Relationships between environmental factors and oscillations in jellyfish abundance, especially in the early life stages, could help to interpret past increases and also predict scenarios in a changing future. For the first time, we present cubozoan spatial and temporal distributions in the earliest stages and their relationships with different factors. Abundances of *Carybdea marsupialis* medusae showed high interannual variability from 2008 to 2014 along the Dénia coast (SE Spain, W Mediterranean). During 2015, samples were collected from 11 beaches along 17 km of coastline, 8 times from January to November in order to determine the effects of environmental factors on the distribution of juvenile *C. marsupialis*. Juveniles (≤ 15 mm diagonal bell width) were present from May to July, with more sampled near shore (0 – 15 m). Most of them occurred in June when their numbers were unequal among beaches (average 0.05 ind m^−3^, maximum 6.71 ind m^−3^). We tested distributions of juveniles over time and space versus temperature, salinity, nutrients (N, P and Si), chlorophyll-*a* (Chl-*a*), and zooplankton abundance. Temperature and cladocerans (zooplankton group) were significantly positively correlated with juvenile distribution, whereas Chl-*a* concentration was weakly negative. By contrast, in 2014, high productivity areas (Chl-*a* and zooplankton) overlapped the maximum adult abundance (5.2 ind m^−3^). The distribution of juveniles during 2015 did not spatially coincide with the areas where ripe adults were located the previous year, suggesting that juveniles drift with the currents upon release from the cubopolyps. Our results yield important insights into the complexity of cubozoan distributions.

## Introduction

The relationships between environmental variables and spatio-temporal distributions of the developmental stages of jellyfish are decisive for revealing their dynamic population and ecological niches. Despite increasing interest in the ecology of jellyfish in recent years due to various factors, such as their negative effects on human activities [1,2] and human health [3], quantitative studies on cubozoan medusae distribution are rare [4]. This is probably because of the infrequent observation of some species, which generally occur in low numbers and show great variation in abundance over a short period of time [4]. There are some spatial studies indicating that polyps and juveniles of *Chironex fleckeri* may inhabit estuaries influenced by freshwater, while adults actively swim to the open ocean, but remain very close to the coastline [5,6]. Between the coastline and the Great Barrier Reef, the abundances of adult cubozoans (*Carybdea xaymacana, Carukia barnesi, Alatina* sp., *Copula sivickisi* and *C. fleckeri*) changed across the shelf on scale of kilometres [7].

*Carybdea marsupialis* (Linneo, 1758), the only box jellyfish species to date described in the Mediterranean Sea, has been regularly observed in high densities in the Adriatic Sea since the 1980s [8,9]. Recent genetic and morphological studies have shown that records referring to *C. marsupialis* in tropical and subtropical regions in the Atlantic Ocean belong to *C. xaymacana* in the Caribbean and *C. branchi* in the South African seas, which suggests that *C. marsupialis* is endemic to the Mediterranean Sea [10].

Along the Spanish Mediterranean coast, *C. marsupialis* adults were first detected in high abundance (maximum 2.65 ind m^−3^) in shallow waters (~1 m depth) off Dénia in the province Alicante during summer 2008 [11]. After 2008, abundances of cubomedusae in this area have shown high inter-annual variability. For instance, the mean density of *C. marsupialis* in 2010-2011 was 0.8 ind m^−3^ [12], but 0.17 ind m^−3^ in 2013 [13]. More recently, this species has been found in Malta [14] and off the east coast of Tunisia, with an average of 1.1 ind m^−3^ and a maximum of 1.8 ind m^−3^ [15].

The seasonality and lifecycle of *C. marsupialis* have been studied in the scope of the LIFE CUBOMED Project on the Spanish Mediterranean coast [12,13,16]. The life cycle of this cubozoan alternates between a planktonic stage as a sexually-reproductive cubomedusa and an asexually-reproductive benthic stage as a cubopolyp. Table 1 shows different stages of C*. marsupialis* according to size, seasonality in Dénia, Alicante, Spain, and sampling techniques.

**Table 1.** Different stages of *Carybdea marsupialis* cubomedusae according to size, their seasonality in Dénia, Alicante, Spain, and sampling technique used for their capture. *DBW: Diagonal Bell Width (equivalent to diameter for small and recently-detached box jellyfish).

| Life cycle | Period | Methods |
| --- | --- | --- |
| Recently-detached DBW*<2 mm | May-July | 500-µm hand-nets and trawling |
| Small 2 ≤ DBW*<5 mm | June-July | 500-µm hand-nets and trawling |
| Medium 5 ≤ DBW*< 15 mm | July-August | 500-µm hand-nets and trawling |
| Adults DBW*≥ 15 mm | August-November | 1-4 mm hand-nets |

The locations of the cubopolyps are still unknown, despite extensive efforts to find them [16]. Manipulative experiments with other cubozoan species have revealed that factors such as salinity, temperature and food supply influence the metamorphosis of cubopolyps into cubomedusae [7,17–19]. These environmental factors are important in determining the life cycle of cubozoans and, therefore, their abundances [4].

Previous analyses of the abundance and distribution of *C. marsupialis* at 0.5-1 m depth off Dénia showed an inverse relationship between medusa abundance and salinity [16] and occurrence of this species in conditions of low salinity, high coastal productivity (Chl-*a* and phosphate) and optimal physical conditions (wind current speed and direction) [12]. Medusa outbreaks also were linked with anthropogenic nutrient inputs (N, P and Si) and high productivity [13]. We built on these earlier works to focus on the distribution of juvenile *C. marsupialis*, by use of a more detailed sampling grid that included points at different depths in the water column and three distances from the coast to overcome the spatial restrictions pointed out by Canepa et al. [12]. Our main objective in 2015 was to determine the relationships between the abundance of juvenile *C. marsupialis* and the environmental variables: temperature, salinity, nutrients (N, P and Si), Chl-*a* and zooplankton. Our null hypothesis was that the distribution of recently-detached *C. marsupialis* medusae was not related to any of those environmental factors, but instead, reflected the spatial distribution of adults in the previous year (2014).

## Materials and methods

### Study area

The study area comprised the coastal waters off Dénia (38° 50’ 33.57’’ N, 0° 6’ 24.83’’ E) in the Western Mediterranean, Spain (Fig 1). Dénia is a coastal tourist town surrounded by intensive irrigated agriculture. It presents a typical Mediterranean climate: rainy springs, mild winters and warm, dry summers, occasionally with storms. Three rivers discharge into the sea along this stretch of coast: the Racons, Almadrava and Alberca rivers. The Racons River is the largest in volume and fed by runoff from citrus and rice croplands and the wastewater treatment plants (WWTP) from the towns of Pego and El Verger North. The wastewaters from the El Verger-Els Poblets WWTP feed into the Almadrava River, while the Dénia-Ondara-Pedreguer WWTP discharges directly into the sea via a marine outfall located some 1300 m off Raset beach [16]. Additionally, some areas along this stretch of coast have aquifers that discharge high-nitrate (~25-100 mg l^−1^) groundwaters into the sea along a narrow strip (tens of metres) close to shore [16].

**Fig 1.**
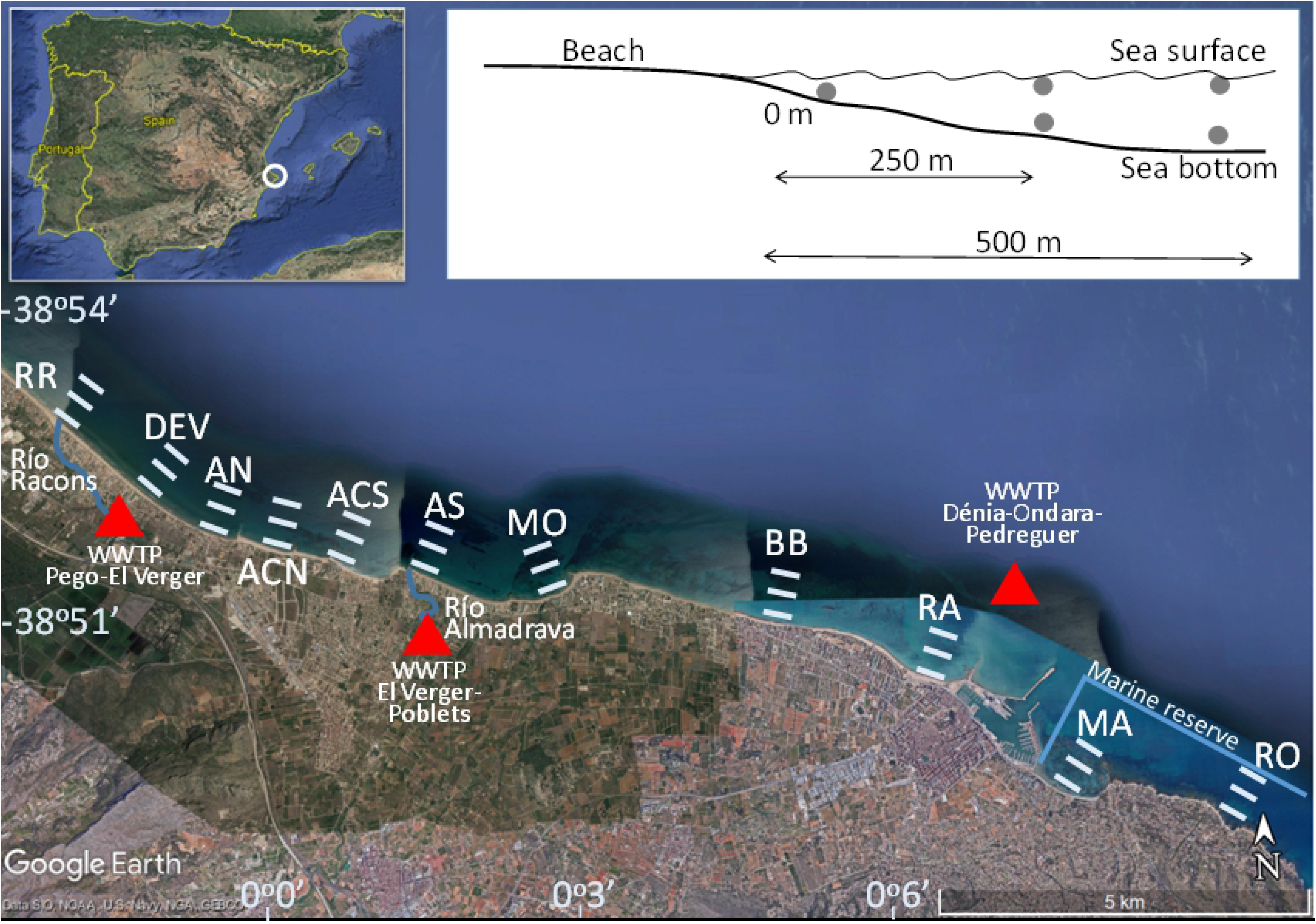
Sampling area for *Carybdea marsupialis* cubomedusae off Spain in the western Mediterranean. Sampling stations (black points): RR: Racons River, DEV: Deveses, AN: Almadrava Norte, ACN: Almadrava Centro-Norte, ACS: Almadrava Centro-Sur, AS: Almadrava Sur, MO: Molins, BB: Blay Beach, RA: Raset, MA: Marineta, RO: Rotas. WWTP: Waste water treatment plant discharge point (triangles).

Samples were taken along 17 km of coastline at 11 beaches (from North to South): Río Racons (RR), Deveses (DEV), Almadrava Norte (AN), Almadrava Centro Norte (ACN), Almadrava Centro Sur (ACS), Almadrava Sur (AS), Molins (MO), Blay Beach (BB), Raset (RA), Marineta Casiana (MA) and Rotas (RO) (Fig 1). All the sampling locations to the north of Dénia harbour (from RR to RA) are long sandy beaches, gently sloping with patchy *Posidonia oceanica* meadows. Specific features characterise certain locations: the waters around RR and DEV are influenced by discharge from the Racons river [16]; AN, ACN, ACS and AS are separated by stone breakwaters built in 2005; both MO and BB are very shallow sandy beaches with rocky patches and *P. oceanica* meadows; RA and MA are on opposite sides of the harbour and somewhat protected, combining sand, mud, and areas of *Caulerpa prolifera*; RO to the south has a rocky shoreline with boulders and pebbles. Both MA and RO are located within the Marine Reserve of Cabo San Antonio.

### Cubomedusae and zooplankton sampling and analysis

The cubomedusae were captured using plankton nets dragged parallel to the coastline. In order to detect the first appearance of new cubomedusae, coastal sampling started in January and used a 500-μm-mesh net to collect them, as suggested by Canepa et al. [12].

In 2015, we took samples from each location in January, March, May, June, July, August, October and November. We sampled at three distances from the coast: within 15 m from the shoreline (hereinafter referred to as “0 m”) walking for 15 min at ~0.4 m s^−1^ and using hand nets (length 1.5 m, mouth area 0.15 m^2^, mesh size 500 μm); at ~250 m and ~500 m using nets with the same dimensions as the hand nets, but towed by a boat moving at ~0.7 m s^−1^ for 5 min.

The nets were equipped with flow meters (KC Denmark Digital and General Oceanics, 2030R) for the subsequent estimation of filtered water volume. At locations where the water was more than 3 m deep, we took two samples, one from the surface and the other from close to the seabed. Densities of adults from the previous year were obtained by the same methodology walking on the shoreline of the beaches with hand nets in September and October 2014 (Fig 2).

**Fig 2.**
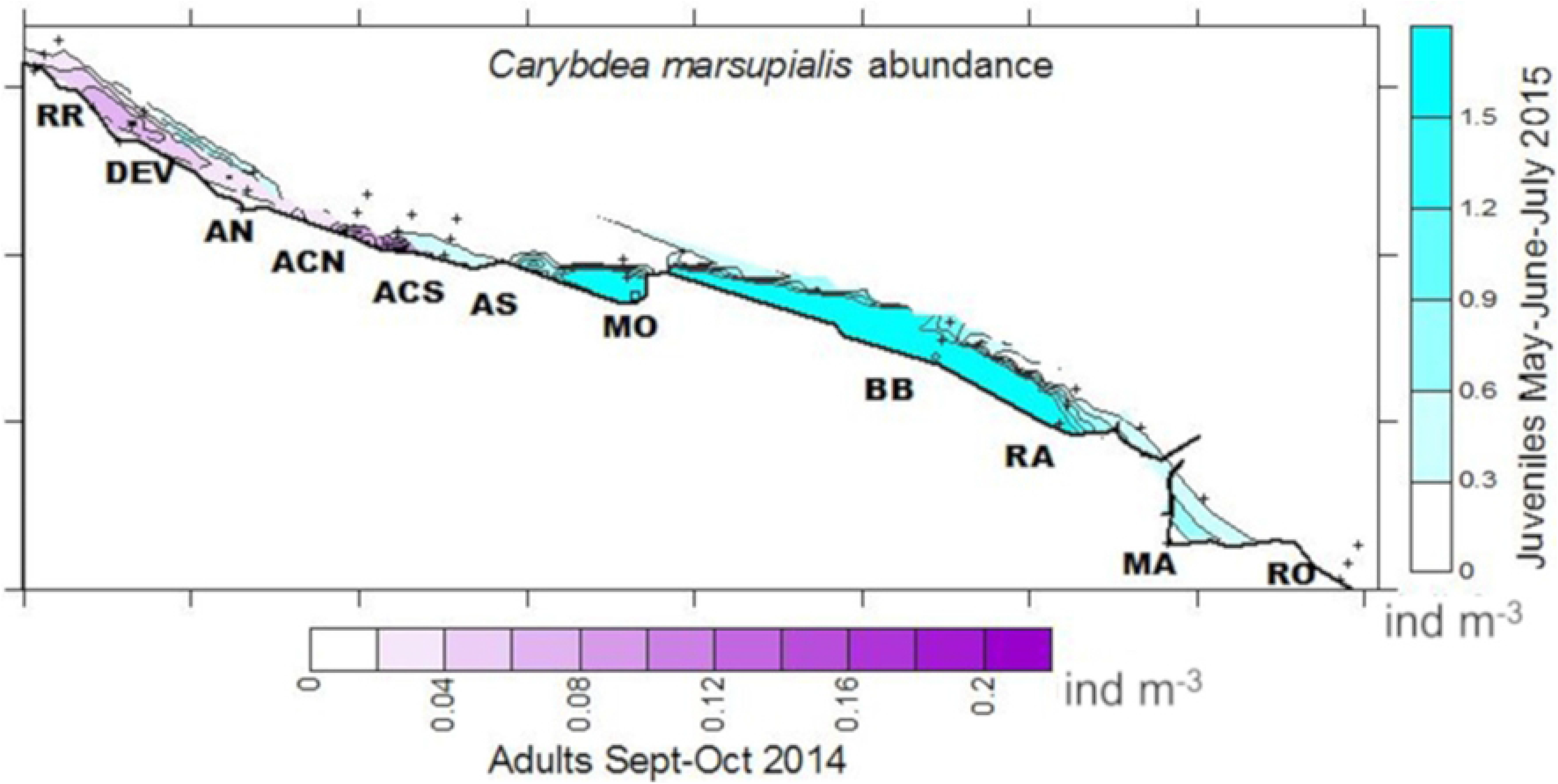
Spatio-temporal distribution of adult cubomedusae in 2014 (present only in September and October) and juvenile abundance in 2015 (present only in May, June and July) (ind m^−3^). Data in both years are from the same sampling locations (abbreviations as in Fig 1) off Spain in the western Mediterranean.

The same procedures were followed for zooplankton as for cubomedusae, except nets with 200-μm mesh size instead of 500-μm were used. The zooplankton and cubozoan nets were towed simultaneously.

Cubomedusa and zooplankton samples were preserved in the laboratory in 4% formalin buffered with sodium tetraborate. The distance between opposite pedalia (diagonal bell width, DBW) was measured for each *C. marsupialis* medusa captured [12,13,16]. An aliquot of each zooplankton sample was taken using a box-type plankton splitter. We counted and measured (total length) a minimum of 150 individuals per sample with the aid of a stereomicroscope (Leica S8APO). Specimens were classified into the following groups: amphipods, cladocerans, copepods, fish eggs and larvae, mollusc larvae, and polychaetes. Counts were standardized to numbers m^−3^.

### Environmental data

At each sampling location, temperature and salinity were measured using a conductivity and temperature logger (CT, JFE Advantech) at 0 m and a conductivity, temperature and depth logger (CTD, Seabird Electronics 19-10) at 250 m and 500 m from the coast. Additionally, two litres of surface and near-bottom water samples were taken at each location for subsequent nutrient (N, P and Si) and Chl-*a* analysis. Water samples were collected by hand with a plastic bottle at the surface and with a Niskin bottle (KC Denmark) at depth. Seawater samples were kept in the dark in an isothermal cooler box with ice packs and transported immediately to the Research Institute for Integrated Management of Coastal Areas of the Polytechnic University of Valencia for analysis. Temperature, salinity, and Chl-*a* were analysed for all the sampling months.

The nutrients from each location were analysed for January, May, July and October. Nutrients (ammonium (NH_4_^+^), nitrite plus nitrate (NO_2_^−^+ NO_3_^-^), phosphate (PO_4_^3^) and silicate (Si(OH)_4_) were analysed according to Aminot & Chaussepied [21]. The precision of these methods was 5% for NH_4_^+^, NO_2_^−^ + NO_3_^−^ and PO_4_^3−^, but 3% for Si(OH)_4_. Dissolved inorganic nitrogen (DIN) was calculated as the sum of NH_4_^+^ and NO_2_^−^ + NO_3_^−^. Chlorophyll *a* was measured using the trichromatic method, based on spectrophotometry following the methodology described in APHA [22].

Surface current vectors were obtained at all sampling locations at 0 m. Surface current vectors also were obtained at RR, ACS, RA and RO at 250 m and 500 m from the shoreline using two drifting buoys at each station and recording their initial and final (5 min after release) waypoints with a GPS device (Garmin 72H) [12,16].

### Statistical analysis

We used the non-parametric Kruskal-Wallis test to examine significant differences in juvenile *C. marsupialis* distributions among months, beaches, distances to the coast and depths in the water column. In addition, a Tukey test (post-hoc pairwise multiple comparisons test) was conducted when Kruskal-Wallis test results were significantly different and more than two groups were present.

To evaluate the effects of environmental variables on the abundances of juvenile *C. marsupialis,* we first used the Pearson correlation test to inspect the association between response variables (i.e. avoid collinearity) [12,23]. Pairs of parameters with a coefficient >0.5 were dropped from the model. The explanatory variables were explored to identify outliers. A transformation (Log_10_) was then applied to Chl-*a*, DIN, phosphate (P), silicate (Si), salinity and temperature (T) variables, because excluding the outliers would result in deficient data. The copepod and cladoceran variables, as well as the response variable *C. marsupialis*, were not log transformed.

Then, generalized linear models (GLM) with Poisson and negative binomial distributions were fitted to the data to explore the association between *C. marsupialis* and environmental parameters for all sampling locations and distances from the coastline [12,23]. The models used a logistic-link function to assess positive fitted values and because of the high variability of the filtered volume (data not shown), this variable was used as an offset in the GLMs [12].

The optimal models were selected using a backward strategy based on the significance of each explanatory variable and the Akaike Information Criterion (AIC). The AIC measures goodness of fit, evaluates the model’s complexity and penalizes an excess of parameters. The lower the AIC, the better the model [23]. Finally, all of the optimal models selected were validated via residual analysis [24].

Statistical analyses were done using the statistical platform R version 3.3.0. Maps showing the surface spatio-temporal distribution of each parameter were generated using SURFER 8.00, using the “triangulation with linear interpolation” method [25].

## Results

### Distribution and abundance of *Carybdea marsupialis*

Abundances of juvenile cubomedusae differed significantly among months (Table 2), with the highest mean density in June (0.26 ind m^−3^; Table 3, Fig 3A). Abundances of *C. marsupialis* juveniles were higher at 0 m (0.18 ind m^−3^) than at 250 m or 500 m (both 0.01 ind m^−3^) (Table 4, Fig 3C). Juvenile abundances were higher at the surface than near the seabed (0.01 vs 0.003 ind m^−3^) (Table 4, Fig 3D). The large number of zero captures in autumn and winter may have caused the lack of significant difference among locations (p-value = 0.06) (Table 2), even though many more juveniles were captured at two sampling stations (MO and BB) than others stations in May and June (Table 3, Fig 3B).

**Fig 3.**
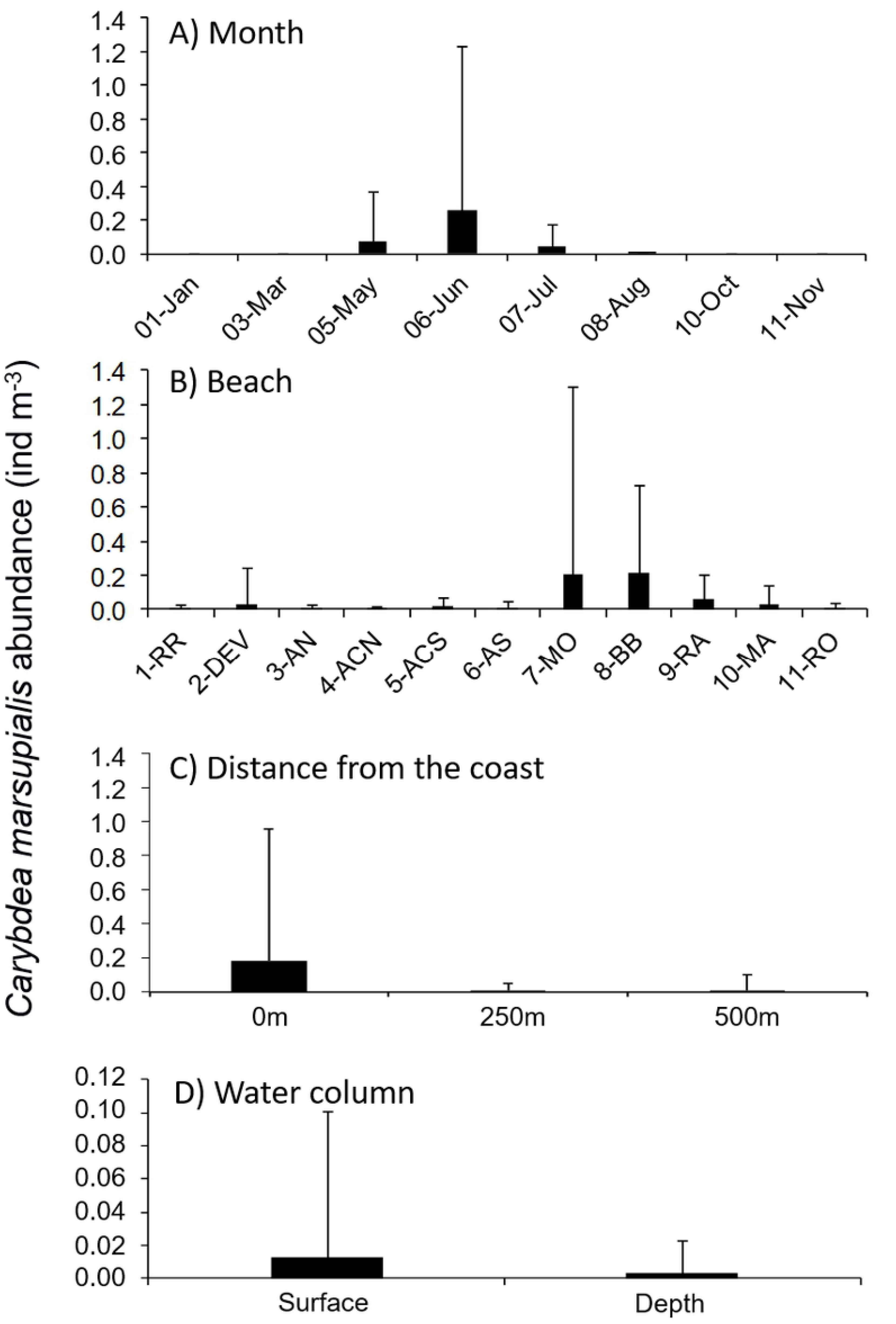
Juvenile *Carybdea marsupialis* abundance in 2015. Mean ±SD of all samples (A) over time, (B) by sampling locations (abbreviations as in Fig 1), (C) by distance from shore (0, 250, 500 m), and (D) by depth in the water column (only stations at 250 and 500 m).

**Table 2.**
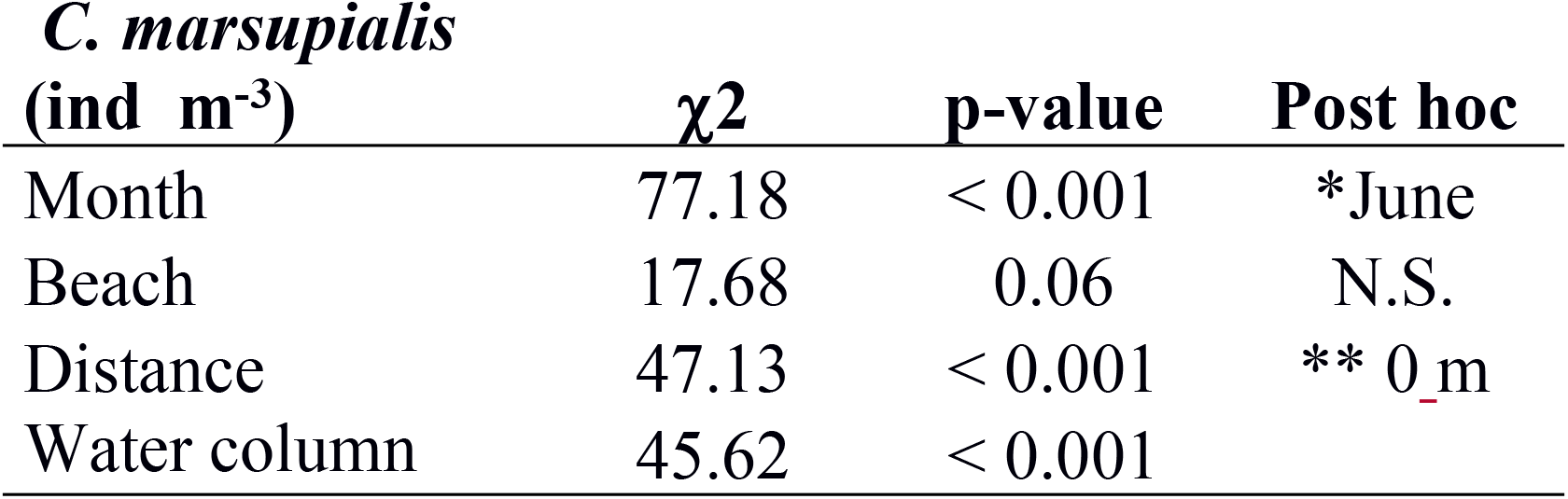
Kruskal-Wallis and Tukey post hoc test results of the variability of juvenile *Carybdea marsupialis* distribution among months, beaches, distances from shore (0, 250, 500 m) and water column (surface, depth). Chi-square: ‘χ^2^’. Significance codes “p-value”: ‘**’ 0.01; ‘*’ 0.05; ‘N.S.’ not significant

**Table 3.** Juvenile *Carybdea marsupialis* abundance (ind m^−3^) (Mean and SD) by month at 11 beaches of Dénia (Alicante, Spain). Station abbreviations as in Fig 1.

|  | January |  | March |  | May |  | June |  | July |  | August |  | October |  | November |  |
| --- | --- | --- | --- | --- | --- | --- | --- | --- | --- | --- | --- | --- | --- | --- | --- | --- |
|  | MEAN | SD | MEAN | SD | MEAN | SD | MEAN | SD | MEAN | SD | MEAN | SD | MEAN | SD | MEAN | SD |
| 1-RR | 0 | 0 | 0 | 0 | 0.004 | 0.009 | 0.030 | 0.067 | 0.010 | 0.022 | 0 | 0 | 0 | 0 | 0 | 0 |
| 2-DEV | 0 | 0 | 0 | 0 | 0 | 0 | 0.264 | 0.590 | 0 | 0 | 0 | 0 | 0 | 0 | 0 | 0 |
| 3-AN | 0 | 0 | 0 | 0 | 0.004 | 0.009 | 0.030 | 0.067 | 0 | 0 | 0 | 0 | 0 | 0 | 0 | 0 |
| 4-ACN | 0 | 0 | 0 | 0 | 0.016 | 0.036 | 0.008 | 0.018 | 0.014 | 0.031 | 0 | 0 | 0 | 0 | 0 | 0 |
| 5-ACS | 0 | 0 | 0 | 0 | 0.062 | 0.139 | 0.030 | 0.067 | 0.018 | 0.025 | 0.004 | 0.009 | 0 | 0 | 0 | 0 |
| 6-AS | 0 | 0 | 0 | 0 | 0 | 0 | 0.088 | 0.088 | 0 | 0 | 0 | 0 | 0 | 0 | 0 | 0 |
| 7-MO | 0 | 0 | 0 | 0 | 0.068 | 0.135 | 1.426 | 2.956 | 0.052 | 0.116 | 0 | 0 | 0 | 0 | 0 | 0 |
| 8-BB | 0 | 0 | 0 | 0 | 0.673 | 1.166 | 0.793 | 0.612 | 0.260 | 0.442 | 0 | 0 | 0 | 0 | 0 | 0 |
| 9-RA | 0 | 0 | 0 | 0 | 0.157 | 0.271 | 0.100 | 0.130 | 0.187 | 0.323 | 0.007 | 0.012 | 0 | 0 | 0 | 0 |
| 10-MA | 0 | 0 | 0 | 0 | 0.070 | 0.121 | 0.145 | 0.290 | 0.053 | 0.086 | 0 | 0 | 0 | 0 | 0 | 0 |
| 11-RO | 0 | 0 | 0 | 0 | 0 | 0 | 0.046 | 0.082 | 0 | 0 | 0 | 0 | 0 | 0 | 0 | 0 |
| ALL SITES | 0 | 0 | 0 | 0 | 0.07 | 0.30 | 0.26 | 0.98 | 0.04 | 0.014 | 0.001 | 0.004 | 0 | 0 | 0 | 0 |

**Table 4.** Average abundances of juvenile *Carybdea marsupialis* cubomedusae at all sampling locations off Spain in the western Mediterranean at different distances from shore (0, 250, 500 m), and depth (surface, deep).

| ***C. marsupialis*** |  |  |  |
| --- | --- | --- | --- |
| **(ind m<sup>-3</sup>)** |  |  |  |
|  | **Distance from**<br>**coastline** | **Mean** | **SD** |
|  | **0 m** | 0.182 | 0.772 |
|  | **250 m** | 0.009 | 0.040 |
|  | **250 m surface** | 0.011 | 0.048 |
|  | **250 m deep** | 0.007 | 0.026 |
|  | **500 m** | 0.008 | 0.091 |
|  | **500 m surface** | 0.013 | 0.116 |
|  | **500 m deep** | 0.000 | 0.000 |
|  | **Surface** | 0.012 | 0.089 |
|  | **Deep** | 0.003 | 0.019 |
|  | **Global** | 0.048 | 0.380 |

Juvenile abundances differed markedly among months. The first recently-detached (< 2 mm DBW) *C. marsupialis* appeared in May (all stations mean 0.07 ind m^−3^), with their peak at station BB at 0 m (2.01 ind m^−3^). Their maximum abundance (6.71 ind m^−3^) was observed in June at station MO at 0 m. Abundances overall were considerably lower in July, a 85% less (0.26 vs. 0.04 ind m^−3^, all stations mean). *C. marsupialis* juveniles were practically non-existent in August 2015, with only two captures (all stations mean 0.001 ind m^−3^) and finally, no juveniles or adults were captured in autumn or winter (Table 3, Fig 3 and 4).

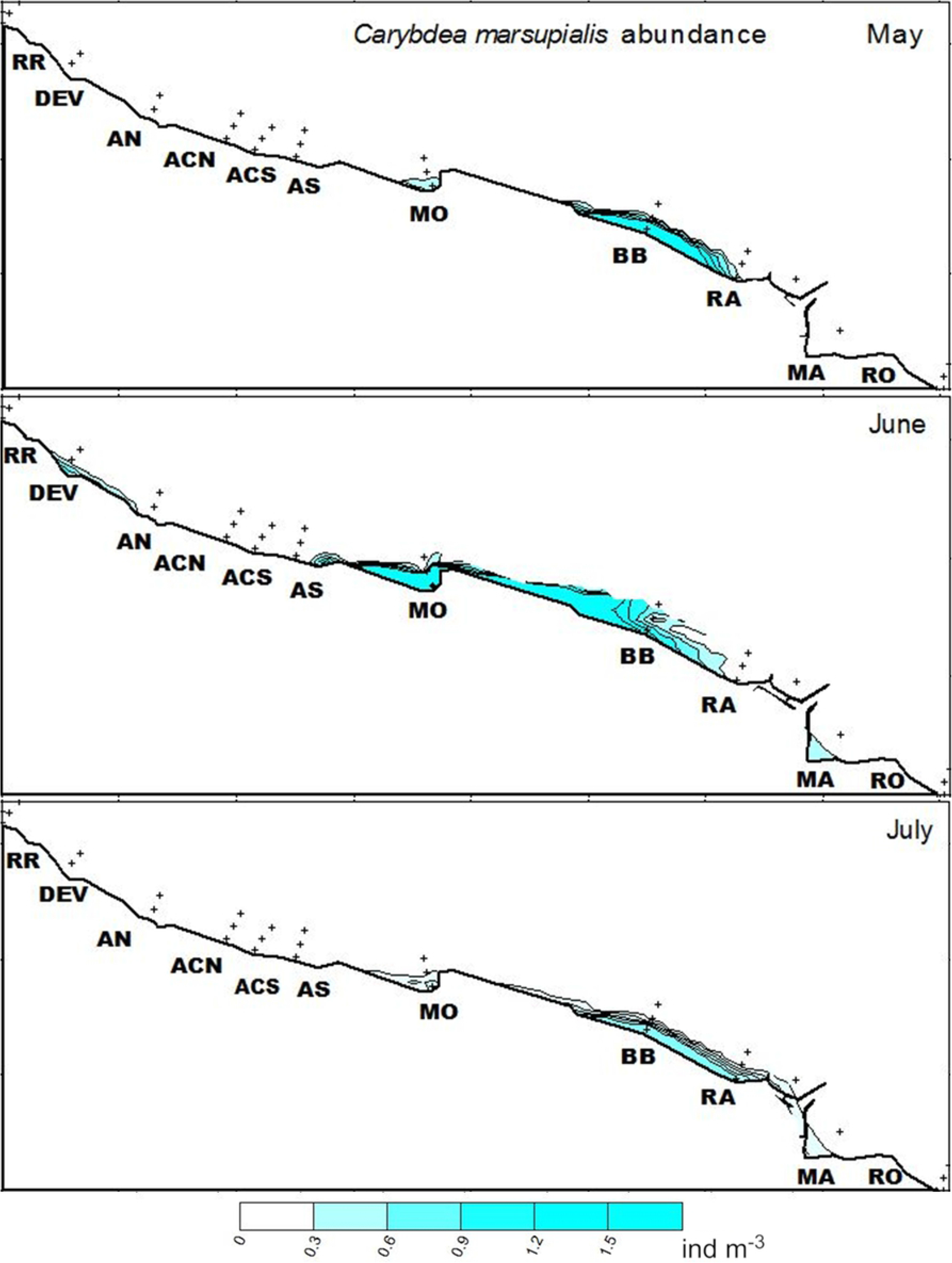

Of the 804 juveniles captured at the shoreline, recently-detached cubomedusae (756 individuals, DBW ≤ 2 mm) were the most numerous of all sizes in May, June and July, (Fig 5). In May, all 153 cubomedusae measured less than 2.2 mm (mean size 1.2 mm). In June, 559 cubomedusae were captured, with a mean DBW of 2.0 mm; the largest one measured 6.2 mm detected at station MA. The widest size distribution was in July (92 individuals with a mean of 2.3 mm), when two of measured 9 and 10 mm at station MA.

**Fig 5.**
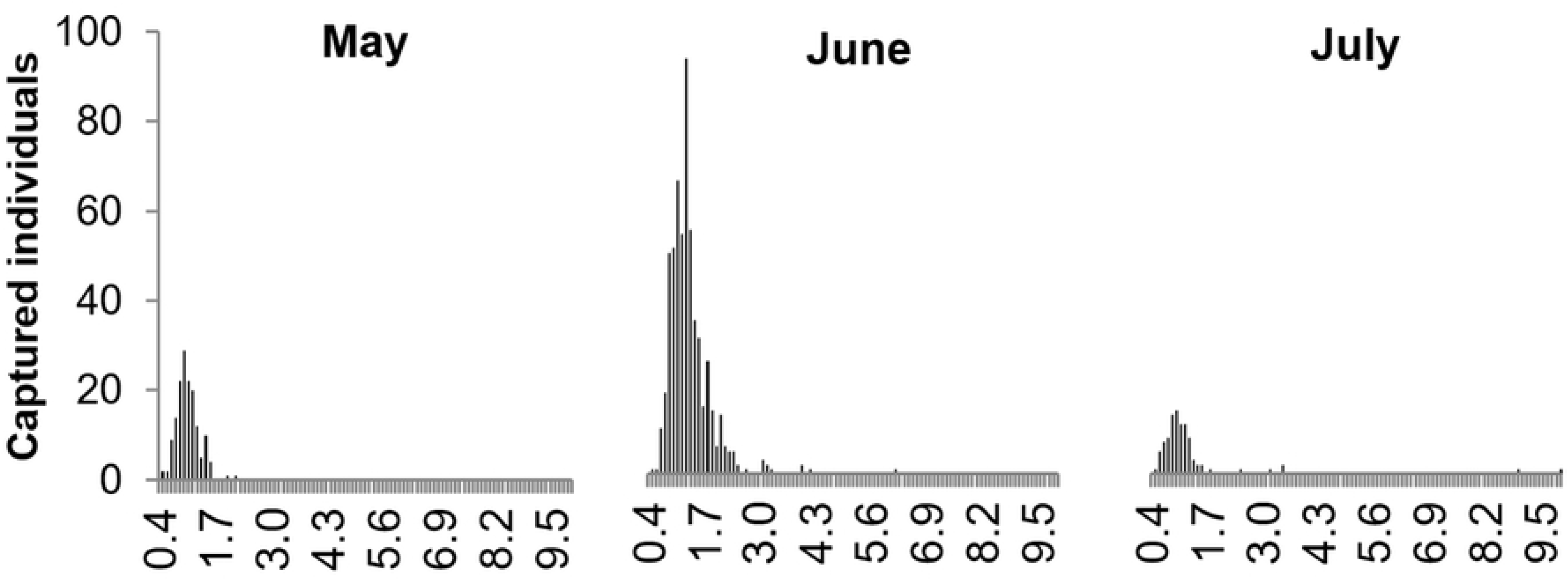
Size frequency distribution of *Carybdea marsupialis* cubomedusae captured at all sampling locations off Spain in the western Mediterranean in May, June and July 2015. Size of individuals is the diagonal bell width, DBW.

### Spatio-temporal variation of environmental variables

The environmental variables followed a typical Mediterranean seasonal pattern. Water temperature was highest in summer (31.3°C at MA at 0 m, where we did not differentiate between surface and deep samples due to the shallowness (~1m) of this area) and lowest in winter (13.1°C at RO at 500 m, deep sample). The first small cubomedusae appeared in May in all stations at 0 m (except DEV, AS and RO), coinciding with the spring increase in temperature; the highest temperature recorded that month was at RA (24.2°C) with the second highest cubomedusa abundance (0.47 ind m^−3^) (S1 Table, S1 Fig). It was in June, however, when *C. marsupialis* abundance peaked with a mean temperature of 23.8°C (all stations); abundances did not increase even though the temperature was higher during the next two months (July 27.9°C and August 27.1°C).

Average salinity for all surface and deep samples was 37.2, with the lowest values found in the northern-most location (RR) in July (17.8) and November (28.1) at 0 m and in January (24.5) at 250 m surface sample, probably due to the Racons River discharge (S2 Table, S2 Fig).

Nutrients and Chl-*a* concentrations had similar seasonal patterns in the sampling area. DIN had low values in January, May and July (3.20, 2.22 and 3.06 μM, respectively), but peaked in October (4.80 μM) (S3 Table, S3 Fig). The P concentrations were low in January and May (mean 0.03 μM), but high in July and October (0.10 and 0.09, respectively) (S4 Table, S4 Fig). May and October had the maximum mean Chl-*a* values of 0.7 and 1.6 μg l^-1^, respectively (S5 Table, S5 Fig).

Concentrations of nutrients and Chl-*a* generally were highest in the north (RR) and second highest near Dénia harbour. Maximum mean values of DIN, ranging from 6.83 to 13.39 μM, were at the northern beach (RR) (Fig S3). Northern beaches, especially RR, had the highest mean P values, reaching 0.72 μM in July (S4 Fig). The spatial pattern of Chl-*a* was less marked than for DIN and P; the northern half of the sampling area generally had higher than average Chl-*a* values. Maximum Chl-*a* were at RR (6.0 μg l^−1^), with the minimum (0.1 μg l^−1^) occurring in many locations in the southern half of the sampling area (S5 Fig).

When juvenile cubomedusae were present in May, June and July, current speeds ranged from 0.5 cm s^−1^ to 22.9 cm s^-1^, averaging 4.1 cm s^−1^ at 0 m, 9.3 cm s^−1^ at 250 m and 11.0 cm s^−1^ at 500 m from the coast. The lowest mean current speed was recorded in May (4.8 cm s^−1^) and similar data were obtained for June (7.0 cm s^−1^) and July (7.5 cm s^−1^). The lowest current speed (0.5 cm s^−1^) recorded for any station was at AS at 0 m, and the highest (22.9 cm s^−1^) was at RO at 500 m.

### Zooplankton community

Zooplankton average abundance was highest during June (3469 ind m^−3^). The northern beaches had the highest abundances, as with Chl-*a*, but zooplankton abundances were lower close to shore than 250 and 500 m off shore. The samples collected in May and October had the fewest zooplankton (479 and 423 ind m^−3^, respectively) (S6 Table, S6 Fig). Zooplankton abundances were similar in surface and deep samples. Copepods, cladocerans and mollusc larvae were most numerous at 250 m from the coastline and were considerably fewer at 0 m.

Copepods were the predominant zooplankton group throughout the study period, representing 67% of the zooplankton total abundance, with an annual average of 733 ind m^−3^. Their maximum density was in June at the northern sampling location RR (20486 ind m^−3^) (S7 Fig). Mollusc larvae were second most abundant (24% of the total abundance), peaking in June (8357 ind m^−3^) and averaging 267 ind m^−3^ among all stations over the year. Cladocerans were third largest group (8% of total abundance), with the highest abundances between May and August (S8 Fig). Conversely, few cladocerans were sampled in January (0.13 ind m^−3^), as compared with the mean density among all months (90.20 ind m^−3^). Cladoceran abundances were highest at all stations in May. Amphipoda, fish eggs and larvae and Polychaeta each represented less than 1% of the total zooplankton density and were not considered further in our analysis. The abundances of copepods, mollusc larvae and cladocerans were higher at the northern stations in the sampling area (DEV and RR) than at other sampling locations. RA had the lowest abundances of these groups.

### Relationships between *C. marsupialis* and environmental factors

The Pearson’s test showed collinearity coefficients > 0.5 between Si concentration and salinity and between total zooplankton and copepods (S7 Table). Consequently, Si and zooplankton were excluded from the model because we considered them less relevant than salinity and copepods as possible explanatory variables for *C. marsupialis* development [12]. The optimal model (GLM Poisson distribution; AIC = 234.08 and dispersion parameter = 0.66) included the interaction between sampling location and distance and the variables selected were salinity, temperature, copepod and cladoceran abundances and concentrations of Chl-*a* and DIN. We used the backward strategy and dropped the P concentration from the model to obtain the lowest AIC value. The model showed a positive significant relationship between temperature and cladoceran abundance and an inverse relationship between Chl-*a* concentration and juvenile *C. marsupialis* abundance. DIN concentration, salinity and copepod abundance did not show significant relationships with *C. marsupialis* distribution (p> 0.05). Stations MO, BB and RA at 0 m distance (p< 0.05) were positively related with *C. marsupialis* distribution, which was negatively related with the distance 250 m from the coastline (Table 5). The explanatory variables Chl-*a*, DIN concentration, copepod and cladoceran density, salinity, temperature, and the interaction between sampling location and distance accounted for 94% of the observed variability.

**Table 5.**
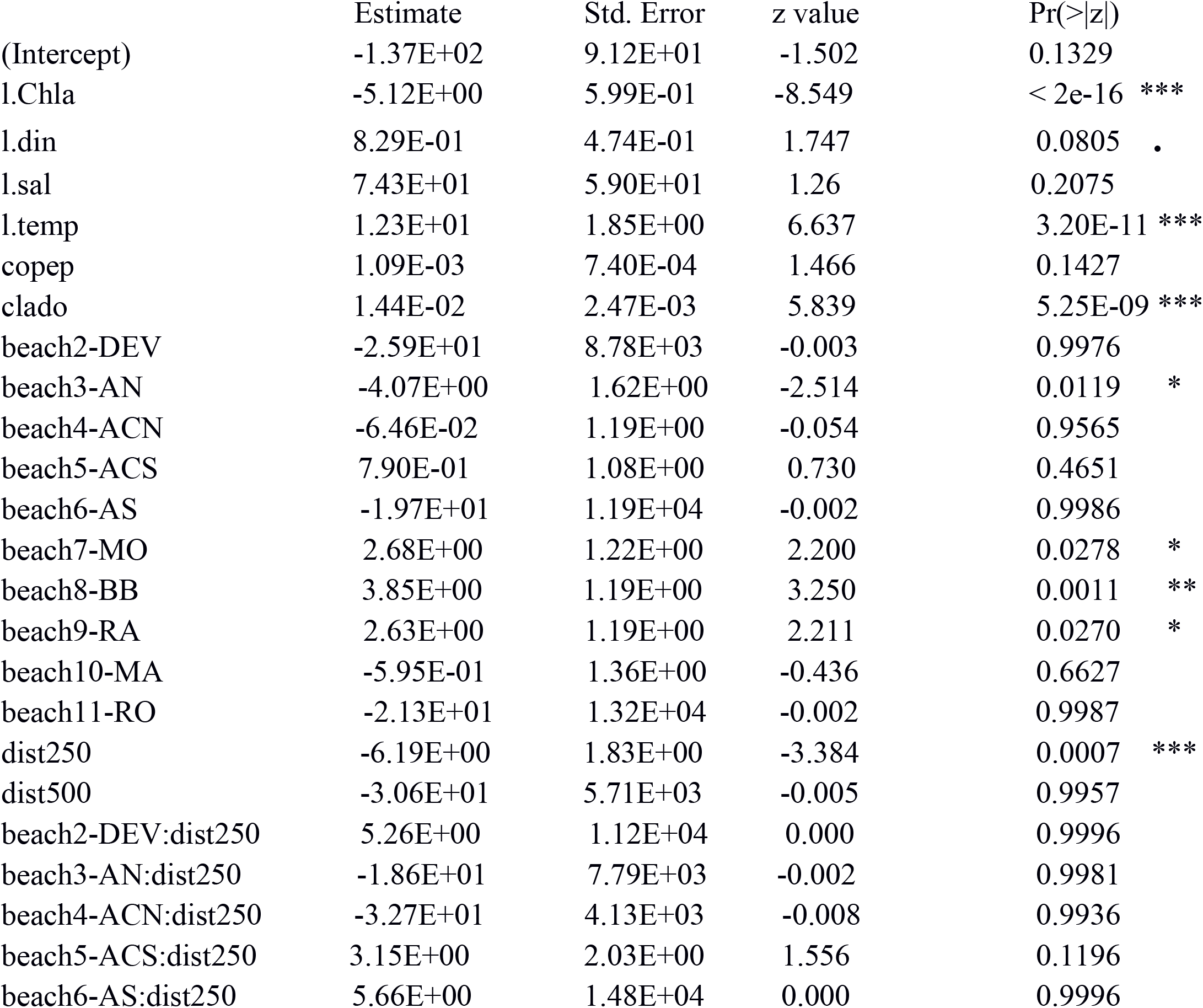
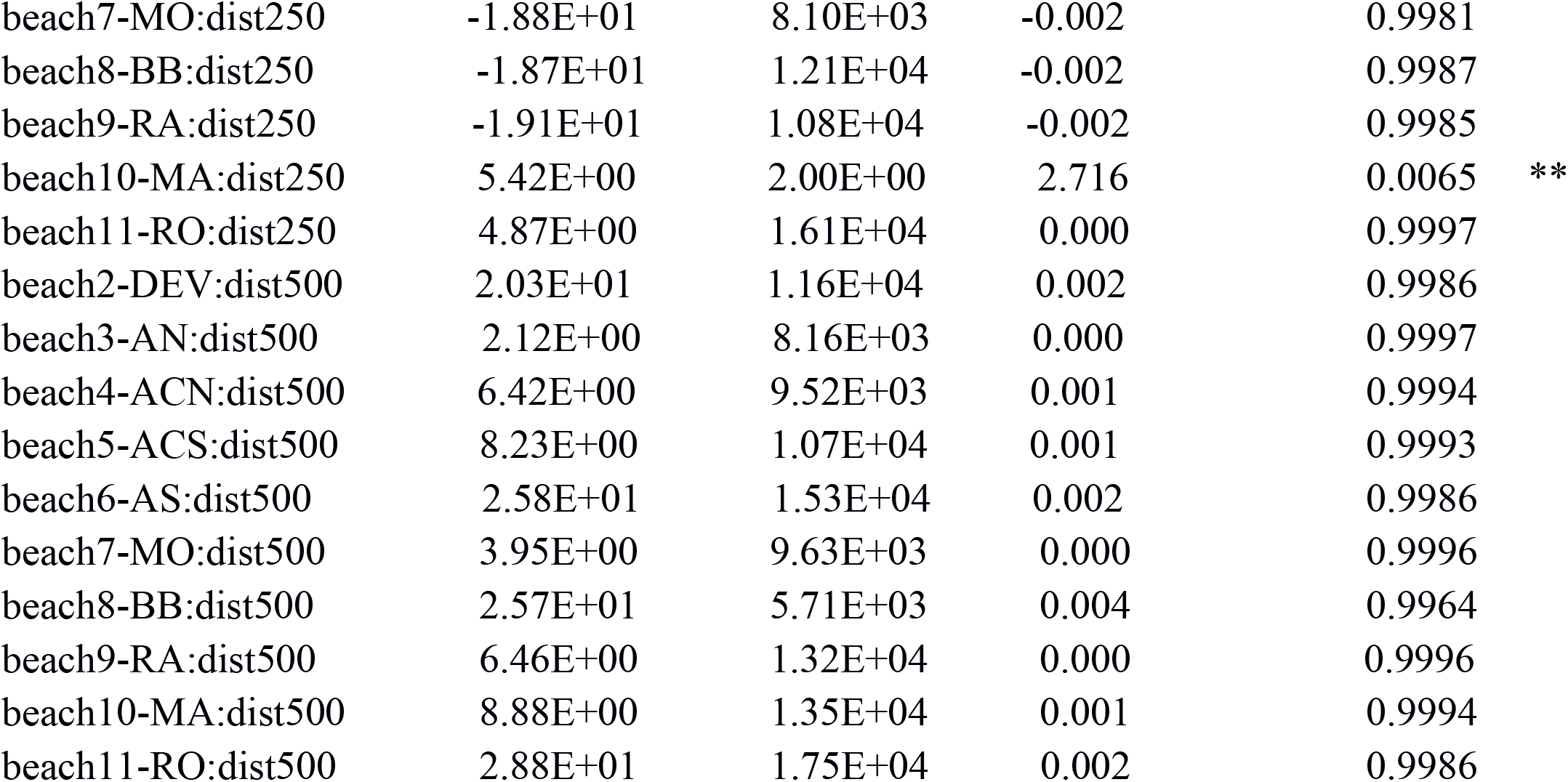
GLM results: relationships between juvenile *Carybdea marsupialis* and chlorophyll-*a* (Chla), dissolved inorganic nitrogen (DIN), salinity (sal), temperature (temp), copepod (copep) and cladoceran (clado) densities and the interaction between beach and distance from the coast (dist250, dist500). Null deviance: 1600.69 on 173 degrees of freedom. Residual deviance: 90.23 on 135 degrees of freedom. Station abbreviations as in Fig 1. Significance codes “z-value”: ‘***’ 0.001; **’ 0.01; ‘*’ 0.05; ‘N.S.’ not significant.

| 344 |  | Estimate | Std. Error | z value | Pr(>\|z\|) |
| --- | --- | --- | --- | --- | --- |
| 345 | (Intercept) | -1.37E+02 | 9.12E+01 | -1.502 | 0.1329 |
| 346 | l.Chl*a* | -5.12E+00 | 5.99E-01 | -8.549 | < 2e-16 *** |
| 347 | l.din | 8.29E-01 | 4.74E-01 | 1.747 | 0.0805 . |
| 348 | l.sal | 7.43E+01 | 5.90E+01 | 1.26 | 0.2075 |
| 349 | l.temp | 1.23E+01 | 1.85E+00 | 6.637 | 3.20E-11 *** |
| 350 | copep | 1.09E-03 | 7.40E-04 | 1.466 | 0.1427 |
| 351 | clado | 1.44E-02 | 2.47E-03 | 5.839 | 5.25E-09 *** |
| 352 | beach2-DEV | -2.59E+01 | 8.78E+03 | -0.003 | 0.9976 |
| 353 | beach3-AN | -4.07E+00 | 1.62E+00 | -2.514 | 0.0119 * |
| 354 | beach4-ACN | -6.46E-02 | 1.19E+00 | -0.054 | 0.9565 |
| 355 | beach5-ACS | 7.90E-01 | 1.08E+00 | 0.730 | 0.4651 |
| 356 | beach6-AS | -1.97E+01 | 1.19E+04 | -0.002 | 0.9986 |
| 357 | beach7-MO | 2.68E+00 | 1.22E+00 | 2.200 | 0.0278 * |
| 358 | beach8-BB | 3.85E+00 | 1.19E+00 | 3.250 | 0.0011 ** |
| 359 | beach9-RA | 2.63E+00 | 1.19E+00 | 2.211 | 0.0270 * |
| 360 | beach10-MA | -5.95E-01 | 1.36E+00 | -0.436 | 0.6627 |
| 361 | beach11-RO | -2.13E+01 | 1.32E+04 | -0.002 | 0.9987 |
| 362 | dist250 | -6.19E+00 | 1.83E+00 | -3.384 | 0.0007 *** |
| 363 | dist500 | -3.06E+01 | 5.71E+03 | -0.005 | 0.9957 |
| 364 | beach2-DEV:dist250 | 5.26E+00 | 1.12E+04 | 0.000 | 0.9996 |
| 365 | beach3-AN:dist250 | -1.86E+01 | 7.79E+03 | -0.002 | 0.9981 |
| 366 | beach4-ACN:dist250 | -3.27E+01 | 4.13E+03 | -0.008 | 0.9936 |
| 367 | beach5-ACS:dist250 | 3.15E+00 | 2.03E+00 | 1.556 | 0.1196 |
| 368 | beach6-AS:dist250 | 5.66E+00 | 1.48E+04 | 0.000 | 0.9996 |

|  |  |  |  |  |  |  |
| --- | --- | --- | --- | --- | --- | --- |
| 369 | beach7-MO:dist250 | -1.88E+01 | 8.10E+03 | -0.002 | 0.9981 |  |
| 370 | beach8-BB:dist250 | -1.87E+01 | 1.21E+04 | -0.002 | 0.9987 |  |
| 371 | beach9-RA:dist250 | -1.91E+01 | 1.08E+04 | -0.002 | 0.9985 |  |
| 372 | beach10-MA:dist250 | 5.42E+00 | 2.00E+00 | 2.716 | 0.0065 | ** |
| 373 | beach11-RO:dist250 | 4.87E+00 | 1.61E+04 | 0.000 | 0.9997 |  |
| 374 | beach2-DEV:dist500 | 2.03E+01 | 1.16E+04 | 0.002 | 0.9986 |  |
| 375 | beach3-AN:dist500 | 2.12E+00 | 8.16E+03 | 0.000 | 0.9997 |  |
| 376 | beach4-ACN:dist500 | 6.42E+00 | 9.52E+03 | 0.001 | 0.9994 |  |
| 377 | beach5-ACS:dist500 | 8.23E+00 | 1.07E+04 | 0.001 | 0.9993 |  |
| 378 | beach6-AS:dist500 | 2.58E+01 | 1.53E+04 | 0.002 | 0.9986 |  |
| 379 | beach7-MO:dist500 | 3.95E+00 | 9.63E+03 | 0.000 | 0.9996 |  |
| 380 | beach8-BB:dist500 | 2.57E+01 | 5.71E+03 | 0.004 | 0.9964 |  |
| 381 | beach9-RA:dist500 | 6.46E+00 | 1.32E+04 | 0.000 | 0.9996 |  |
| 382 | beach10-MA:dist500 | 8.88E+00 | 1.35E+04 | 0.001 | 0.9994 |  |
| 383 | beach11-RO:dist500 | 2.88E+01 | 1.75E+04 | 0.002 | 0.9986 |  |
| 384 |  |  |  |  |  |  |

## Discussion

Temperature and cladoceran abundance were the most influential explanatory variables for the distribution of *C. marsupialis* juveniles, which agrees with earlier reports. Temperature was the main environmental factor affecting the spatio-temporal abundance of all sizes of *C. marsupialis* along the SW Mediterranean coast as found by Canepa et al. [12] and the interannual variability of cladoceran abundance was strongly correlated with the distribution of cubomedusae as described by Acevedo [13].

In 2015 the water temperature increased dramatically from March to May from 14.5°C to 20.3°C (S1 Fig), as typical for the seasonal pattern of the Mediterranean climate. This increase in temperature could be crucial in the life cycle of the species and may have triggered cubopolyp metamorphosis, as found for other species of Cubozoa [4], and led to the first occurrence of small cubomedusae in May. This temporal pattern, with the first recently-detached *C. marsupialis* appearing in May (Fig 4), matched the results for the same area in previous years [12,13,16]. Therefore, we believe that this seasonality is characteristic for the species.

One-day-old *Carybdea morandinii* cubomedusae ~0.5 mm in bell diameter with pedalia appeared only at 15-20 days after liberation from the polyps in the laboratory [26]. Therefore, the small size of the first cubomedusae (mean 1.2 mm diameter) we detected in May indicated they were recently detached medusae. If both species develop similarly, the first cubomedusae collected in Dénia, most of them (78 individuals of a total of 153) with DBW < 1mm and visible small tentacles could be about two weeks after liberation. Furthermore, we also found recently-detached cubomedusae (DBW< 2 mm) in June and July, which suggests that metamorphosis could be a semi-continuous event [12,13] with high interannual variation, e.g. for 2009, recently-detached juveniles were five times more abundant in July than in June (~0.2 vs. ~1.0 ind m^−3^)[16].

This temporal pattern in *C. marsupialis* juvenile abundance could reflect an optimal temperature for metamorphosis in June. Further temperature increase coincided with fewer juveniles, as also observed by Canepa et al. [12]). Experimental evidence connecting the rate of metamorphosis and juvenile mortality at high temperatures remains to be gathered. In addition, three months of metamorphosis may have exhausted the supply of polyps, because *Carybdea* spp. polyps are formed from planulae from adult sexual reproduction the previous summer, which then metamorphose into juveniles the following spring [17]. This suggests that the low abundance of juveniles in 2015 could be related not only to different environmental conditions, but also to fewer new polyps generated at the end of 2014.

Since what it is written in Bordehore 2014, the differences in salinity among the sampling locations were due to the presence of the rivers (mainly at RR) and the discharge of groundwater into the sea (quantified in ~126 10^6^ m^3^ y^−1^ and detected at RR and secondarily at DEV, AS, MO and MA, S2 Fig). Bordehore [16] and Canepa et al. [12] found that abundances of adults and of juveniles plus adults, respectively, were negatively related to salinity. We found, however, that the abundance of juvenile cubomedusae in 2015 was not significantly correlated with salinity (p-value > 0.1) (Table 5). This spatial distribution of juveniles might be due to a combination of active swimming, polyp location and current advection, with the latter two being the most important factors, as we discuss later.

The rate of metamorphosis of *Carybdea* sp. cubopolyps from the Caribbean was higher at the lowest tested salinity (32) than at 35 or 38 in laboratory experiments [19]; those low salinity conditions probably would occur in sheltered bays or estuaries with freshwater inflow. In our field study, all measured salinities ranged from 36.6-38.2 (surface and deep samples included) for May and June, with the exception of sampling location RR (min. of 34.7) where levels are affected by the Racons River (S2 Fig), but where the abundance of *C. marsupialis* juveniles was low (Fig 4). The lowest salinity value in this location was registered in July (17.8) when a unique small cubomedusae was captured, and also in November (28.7) without captures, both at 0 m. The distribution of *C. marsupialis* juveniles was not centred at the low-salinity station (RR). Experiments similar to those by Canepa et al. [19] should be conducted for *C. marsupialis* to test its metamorphosis rate versus salinity.

Therefore, the distribution of *C. marsupialis* seems to be influenced by a combination of environmental factors, with different responses depending on the stage of the life cycle. In this study, we focused on the juvenile stage of cubomedusae. The GLM results did not indicate that nutrients affected their distribution. Phosphorous and DIN concentration were not significantly correlated with *C. marsupialis* abundances (Table 5) in the earliest medusa stages, which could be more affected by dispersion related variables (i.e. wind and current speed and direction), as indicated by Canepa et al. [12].

Nevertheless, earlier studies in the sampling area indicated positive relationships between nutrient concentrations and abundances of all stages of *C. marsupialis* [13], which suggested that as well as contributing to eutrophication in coastal waters [16], nutrient inputs also may lead to a positive bottom-up effect on *C. marsupialis* abundance. Previously, Bordehore [16] estimated percentages of nutrients for the sampling area, mainly nitrates and phosphates from anthropogenic activities and discharged by the Racons River and the wastewater treatment plants. In addition, Acevedo [13] found a negative relationship between nitrate concentration and salinity, as also shown in our results (S7 Table). The stations with elevated DIN and P (S3 and S4 Figs) show the effects of human activities, such as agriculture and wastewater, on eutrophication. Canepa et al. [12] also determined that high abundances of *C*. *marsupialis* medusae (all sizes) were associated with variables indicating high local productivity along the coast of Dénia. Although our results from 2015 did not show a positive relationship between small *C. marsupialis* and Chl-*a* concentration, it is noteworthy that Chl-*a*, as a proxy for primary production, registered peaks in stations with low salinity and high nutrient concentrations, mainly station RR (S5 Fig).

The spatial pattern of *C. marsupialis* juveniles across the shelf showed that their abundance diminished rapidly as distance from the shoreline increased over a few hundred meters. Similarly, in NE Australia, Kingsford et al. [7] showed that abundances of adult cubomedusae of different species were much lower on the outer coral reef than in the inner and mid reef, a strip of ~80 km.

Conversely, zooplankton was more abundant at 250 and 500 m than at 0 m from the shore (S6 Fig). High abundances of *C. marsupialis* were found at 0 m for beaches with relatively low zooplankton levels; however, considering the ingestion rate of this species [13] and the difference in abundance between *C. marsupialis* and zooplankton (~1:10000, Figs 3 and S7), we discounted a potential for top-down control by the cubomedusae.

Our GLM analysis showed a positive relationship between cladoceran abundance and small *C. marsupialis* (Table 4); however, copepod abundance was not significantly correlated with small cubomedusae. Cladocerans peaked in May (S8 Fig), coinciding with the first captured juvenile cubomedusae. As shown by Acevedo [13], release of juveniles and the peak of cladocerans could be linked. In laboratory experiments, *C. marsupialis* revealed an ontogenetic shift in preferred prey, with younger stages being more dependent on cladocerans than adults and she concluded that the survival of smaller stages was related to the abundance of cladocerans [13].

It has been widely demonstrated that primary production plays a significant role in the abundance of zooplankton [27], which could affect cubozoan and scyphozoan distributions. For example, *Alatina moseri* cubomedusae were positively correlated with both factors in Hawaii [28], and Hamner et al. [29] found vertical migrations of *Aurelia aurita* to track their copepod prey.

Optimal development of *C. marsupialis* depends on zooplankton availability, complex interrelations between environmental parameters, and on a combination of physical factors including currents, morphology of the coast, type of substrate and the presence of artificial breakwaters, which could act in retention. Therefore, *C. marsupialis* distribution relies a great deal on the littoral dynamics of the sampling area as well as on the swimming behaviour of the species in different stages.

Within the framework of the LIFE CUBOMED project, experiments on swimming behaviour were conducted in controlled conditions to investigate which sizes of *C. marsupialis* could overcome currents. The average swimming speed of *C. marsupialis* juveniles we recorded in the laboratory ranged between 0.5 and 2.7 cm s^−1^ (average 1.3 cm s^−1^) (S9 Fig), which were similar to those obtained in studies with cubomedusae *Chiropsella bronzie* and *Chironex fleckeri* [30,31]. Current speeds measured in our study area at 0 m averaged 5.8 cm s^−1^ in May, 3.9 cm s^−1^ in June, and 4.9 cm s^−1^ in July (Fonfría et al. unpublished data). Due to the weak swimming ability of juveniles, marine currents would represent the most important influence on small cubomedusa dispersion. Considering the seasonality of *C. marsupialis* in our sampling area and the data obtained from drifting buoys (in 2015 and also 2013-2014, which include several specific factors such as hourly and weather conditions), we calculated that 19.3% of juveniles and 64.0% of adults could overcome currents at 0 m. Previous studies on currents suggest that small juveniles passively drift with the currents, which would play an essential role in dispersion of the species [12]. Conversely, adult *C. marsupialis* medusae would be able to overcome local currents, they present a complex swimming behaviour related with the development of their rhopalia [32–34].

We found the maximum abundances of juvenile cubomedusae in the central stations of the study area, far from the maximum values of Chl-*a* and zooplankton and marine construction. Although the small juveniles have weak swimming capacity, the strong swimming ability of adults lets them select their habitat [35]. Currently, a modelling study is being conducted by the authors in order to understand the movements of *C. marsupialis* juveniles in the study area.

Some authors point out that the increase of artificial constructions along the Mediterranean coast (harbours, breakwaters) may have increased the available habitat for *C. marsupialis* since that could have encouraged the settlement of cubomedusa polyps [11,15,36]. Despite our efforts to locate polyps of *C. marsupialis* by inspection of hard substrates and by using inverse traps, we were unable to find any in our study area. Therefore, the environmental conditions required for polyp development remain unknown.

We assume that polyps were present in the study area because of the detection of recently-detached cubomedusae (DBW≤ 2 mm). The size structure of the population showed that specimens 0.5-1 mm DBW continually appeared throughout May, June and July, where 1 mm DBW was the most frequent size. This suggests that metamorphosis of cubopolyps is not a single event, but rather that cubomedusae are released in different cohorts, in agreement with Acevedo [13].We hypothesized that cubopolyps might be located where the adults mated the previous year. The negative buoyancy of the planulae generated after mating (Bordehore pers. obs.) means that they must sink and settle on the surrounding substratum, as Hartwick [5] described for *C. fleckeri*; however, we found no spatial correlation between areas with maximum abundances of adults and juveniles the following year. We hypothesize this may reflect the current advection of small juveniles (Fig 4).

As explained by the supply-side ecology [37], the sustainability of marine species with benthic and planktonic stages, especially along open coastlines, depends upon recruitment success generation after generation, which is affected by propagule production by mature adults coming from any population (local or upstream), current advection and swimming capabilities of new planktonic individuals. Our understanding of this benthic and planktonic coupling is crucial [38] to understand phenomena such as a cubozoan outbreak [11] and changing abundances among generations. The studied local population of *C. marsupialis* seems to be maintained by recruitment year after year, probably due to a favourable current pattern that retains the recently-detached juveniles. The next step would be to model the advection of juveniles, while considering their swimming capabilities, to ascertain whether this population is maintained in the area due to physical retention by external factors or through the swimming behaviour of individuals that reduces exportation.

Additionally, *C. marsupialis* suddenly disappeared from the area in August 2015, when we captured just two individuals (1.5 and 3.2 mm DBW), compared to other years when adults (>15 mm DBW) were present during August to October (Fig 2). Growth to adult sizes (15 ≤ DBW ≤ 35 mm) was obvious each year from 2008 to 2014 [12,13,16,39], with abundances diminishing as size increased, presumably due to both mortality and current advection, which becomes less relevant as size and swimming ability increase [16]. The authors interpreted that their disappearance could constitute an adult recruitment failure, whose possible causes were not obvious, mainly because year 2015 had similar environmental conditions as in previous years. Higher juvenile mortality, current advection, or a combination of both is one possibility. Severe storms decreased the adult *C. marsupialis* population from 0.2 ind m^−3^ to 0.007 ind m^−3^ in less than one week in 2009 [16]; however, no significant storms occurred in the sampling area during August 2015.

## Conclusion

Temperature, cladoceran abundance, and distance to the coast had positive relationships with juvenile *Carybdea marsupialis* distribution, while Chl-*a,* had a negative relationship (Table 5). The negative relationship between Chl-*a* and small cubomedusae was opposite that of the adults observed in previous years, which would indicate ontogenetic differences in the relationships with environmental variables. Areas of high productivity (Chl-*a* and zooplankton) did not overlap with areas of maximum abundance of small cubomedusae probably because currents were important for juvenile dispersion. The distribution of *C. marsupialis* juveniles during 2015 did not spatially coincide with the areas where ripe adults were located the previous year, supposedly mated and released planulae. Our results suggest that juveniles drift with the currents upon release from the cubopolyps. More studies are needed to establish the location of the polyps and the environmental conditions needed for their development.

## Acknowledgements

We are grateful for the collaboration of *Fundació Baleària*, *Marina El Portet de Dénia* and *Marina de Dénia*. This work is a contribution from the “Ramon Margalef” Environmental Research Institute (IMEM) from the University of Alicante, Spain. We are especially grateful for the sampling and laboratory support provided in the Montgó Research Station by NGO ACIF Marina Alta volunteers: Ángela Alba, Ainara Ballesteros, Miguel Escolano, Ángel Fernández, Marta Gil, Héctor Gutiérrez, Ainara Lacalle and Alba Pérez. We would also like to extend our thanks to Jordi Alventosa and Júlia Escrivá from IGIC-Polytechnic Valencia University for their contribution to nutrient analysis. Editing services were provided by Sea Pen Scientific Writing.

## Supporting Information

S1 Fig. Spatio-temporal distribution of temperature in sampling area during 2015. From north to south: Río Racons (RR), Deveses (DEV), Almadrava Norte (AN), Almadrava Centro Norte (ACN), Almadrava Centro Sur (ACS), Almadrava Sur (AS), Molins (MO), Blay Beach (BB), Raset (RA), Marineta Casiana (MA) and Rotas (RO).

S2 Fig. Spatio-temporal distribution of salinity in sampling area during 2015. From north to south: Río Racons (RR), Deveses (DEV), Almadrava Norte (AN), Almadrava Centro Norte (ACN), Almadrava Centro Sur (ACS), Almadrava Sur (AS), Molins (MO), Blay Beach (BB), Raset (RA), Marineta Casiana (MA) and Rotas (RO).

S3 Fig. Spatio-temporal distribution of DIN concentration in sampling area during 2015. From north to south: Río Racons (RR), Deveses (DEV), Almadrava Norte (AN), Almadrava Centro Norte (ACN), Almadrava Centro Sur (ACS), Almadrava Sur (AS), Molins (MO), Blay Beach (BB), Raset (RA), Marineta Casiana (MA) and Rotas (RO).

S4 Fig. Spatio-temporal distribution of P concentration in sampling area during 2015. From north to south: Río Racons (RR), Deveses (DEV), Almadrava Norte (AN), Almadrava Centro Norte (ACN), Almadrava Centro Sur (ACS), Almadrava Sur (AS), Molins (MO), Blay Beach (BB), Raset (RA), Marineta Casiana (MA) and Rotas (RO).

S5 Fig. Spatio-temporal distribution of Chl-*a* concentration in sampling area during 2015. From north to south: Río Racons (RR), Deveses (DEV), Almadrava Norte (AN), Almadrava Centro Norte (ACN), Almadrava Centro Sur (ACS), Almadrava Sur (AS), Molins (MO), Blay Beach (BB), Raset (RA), Marineta Casiana (MA) and Rotas (RO).

S6 Fig. Spatio-temporal distribution of zooplankton density in sampling area during 2015. From north to south: Río Racons (RR), Deveses (DEV), Almadrava Norte (AN), Almadrava Centro Norte (ACN), Almadrava Centro Sur (ACS), Almadrava Sur (AS), Molins (MO), Blay Beach (BB), Raset (RA), Marineta Casiana (MA) and Rotas (RO).

S7 Fig. Variability of copepod density among months and beaches. Mean ± SD.

S8 Fig. Variability of cladoceran density among months and beaches. Mean ± SD.

S9 Fig. Average swimming speed of *Carybdea marsupialis* juveniles according to size (DBW: Diagonal Bell Width) recorded in the laboratory (Bordehore et al, in prep).

S1 Table. Temperature (Mean and SD) by month at 11 beaches of Dénia (Alicante, Spain). From North to South: Río Racons (RR), Deveses (DEV), Almadrava Norte (AN), Almadrava Centro Norte (ACN), Almadrava Centro Sur (ACS), Almadrava Sur (AS), Molins (MO), Blay Beach (BB), Raset (RA), Marineta Casiana (MA) and Rotas (RO).

S2 Table. Salinity (Mean and SD) at each beach of Dénia (Alicante, Spain), by month. From North to South: Río Racons (RR), Deveses (DEV), Almadrava Norte (AN), Almadrava Centro Norte (ACN), Almadrava Centro Sur (ACS), Almadrava Sur (AS), Molins (MO), Blay Beach (BB), Raset (RA), Marineta Casiana (MA) and Rotas (RO).

S3 Table. Summary of the dissolved inorganic nitrogen (DIN) (Mean and SD) at each beach of Dénia (Alicante, Spain), by month. From North to South: Río Racons (RR), Deveses (DEV), Almadrava Norte (AN), Almadrava Centro Norte (ACN), Almadrava Centro Sur (ACS), Almadrava Sur (AS), Molins (MO), Blay Beach (BB), Raset (RA), Marineta Casiana (MA) and Rotas (RO).

S4 Table. Summary of the Phosphorus (Mean and SD) at each beach of Dénia (Alicante, Spain), by month. From North to South: Río Racons (RR), Deveses (DEV), Almadrava Norte (AN), Almadrava Centro Norte (ACN), Almadrava Centro Sur (ACS), Almadrava Sur (AS), Molins (MO), Blay Beach (BB), Raset (RA), Marineta Casiana (MA) and Rotas (RO).

S5 Table. Summary of chlorophyll-*a* (Mean and SD) at each beach of Dénia (Alicante, Spain), by month. From North to South: Río Racons (RR), Deveses (DEV), Almadrava Norte (AN), Almadrava Centro Norte (ACN), Almadrava Centro Sur (ACS), Almadrava Sur (AS), Molins (MO), Blay Beach (BB), Raset (RA), Marineta Casiana (MA) and Rotas (RO).

S6 Table. Summary of zooplankton abundance (Mean and SD) at each beach of Dénia (Alicante, Spain), by month. From North to South: Río Racons (RR), Deveses (DEV), Almadrava Norte (AN), Almadrava Centro Norte (ACN), Almadrava Centro Sur (ACS), Almadrava Sur (AS), Molins (MO), Blay Beach (BB), Raset (RA), Marineta Casiana (MA) and Rotas (RO).

S7 Table. Collinearity analysis among environmental variables.

